# Inhibition of epigenetic and cell cycle-related targets in glioblastoma cell lines: onametostat reduces proliferation and viability in both normoxic and hypoxic conditions

**DOI:** 10.1101/2023.10.28.564517

**Authors:** Darja Lavogina, Mattias Kaspar Krõlov, Hans Vellama, Vijayachitra Modhukur, Valentina Di Nisio, Helen Lust, Kattri-Liis Eskla, Andres Salumets, Jana Jaal

## Abstract

The choice of targeted therapies for treatment of glioblastoma patients is currently limited, and most glioblastoma patients die from the disease recurrence. Thus, systematic studies in simplified model systems are required to pinpoint the choice of targets for further exploration in clinical settings. Here, we report screening of 5 compounds targeting epigenetic writers or erasers and 6 compounds targeting cell cycle-regulating protein kinases against 3 glioblastoma cell lines following incubation under normoxic or hypoxic conditions. The viability assay indicated that PRMT5 inhibitor onametostat was endowed with high potency under both normoxic and hypoxic conditions in both MGMT-positive and MGMT-negative cell lines. In U-251 MG and U-87 MG cells, onametostat also affected the spheroid formation at concentrations lower than the currently used chemotherapeutic drug lomustine. Furthermore, in T98-G cell line, treatment with onametostat led to dramatic changes in the transcriptome profile by inducing the cell cycle arrest, suppressing RNA splicing, and down-regulating several major glioblastoma cell survival pathways. In this way, we confirmed that inhibition of epigenetic targets might represent a viable strategy for glioblastoma treatment even in the case of decreased chemo- and radiation sensitivity, although further studies in clinically more relevant models are required.

## 1. Introduction

Glioblastoma (GB) is a highly aggressive and lethal form of brain cancer in adults. Despite the use of multiple treatment strategies, the prognosis remains poor. For nearly 2 decades, the standard treatment of patients with GB consists of surgery followed by radiochemotherapy that results in median overall survival time only up to 14.6 months ^1^. Unfortunately, almost all GB patients experience rapid tumour recurrence, and there is no established standard of care for disease progression after radiotherapy and concomitant temozolomide. Lomustine, a nitrosourea compound, is commonly used in chemotherapy when the disease recurs, based on several phase II and III studies that have reported median overall survival times ranging from 5.6 to 9.8 months ^2–4^. Re-irradiation has also been shown to provide temporary disease control in recurrent GB patients ^5^. Since GB is one of the most angiogenic tumours, angiogenesis inhibitors have also been widely tested. Out of these, vascular endothelial growth factor (VEGF) or its receptor (VEGFR) blockers, such as bevacizumab and regorafenib, have shown only modest efficacy ^6,7^.

As radiotherapy, chemotherapy, and angiogenesis inhibitors only result in temporary disease stabilization, additional, more targeted strategies for treating GB are needed. Although GB is a highly vascularized tumour, it is known that blood flow in these newly formed blood vessels is rather inefficient – which leads to the formation of large hypoxic areas and associated necrosis, one of the hallmarks of GB morphological diagnosis ^8^. Therefore, from clinical point of view, the optimal new drug should be equally effective in oxygenated as well as in hypoxic tumour areas.

While numerous reports have investigated the efficiency of individual compounds in simplified model systems such as GB cell lines, few studies have focused on the systematic comparison of different pathways that define the characteristic viability profile of GB. Moreover, few publications have addressed in this context the importance of hypoxia – although it has been shown that the hypoxia-induced pathways affect metabolism, survival, and drug resistance mechanisms of GB cell lines, making hypoxic conditions relevant for screening putative drug candidates ^9–11^.

In a previous study, we screened 13 individual compounds and combinations thereof with temozolomide and lomustine using viability assay for the assessment of targeted modulation in GB cells ^12^. The efficacy of targeted modulation of signalling pathways was compared in two GB cell lines, U-251 MG and T98-G. These cell lines differ in expression levels of O6-methylguanine-DNA methyltransferase (MGMT), which contributes to the resistance of cells to drugs that act via DNA alkylation. We found that among the choice of screened compounds, the inhibitors of Aurora family possessed the highest efficacy. In case of compound combinations, mixtures containing 5-azacytidine – an inhibitor of DNA methyltransferases (DNMT) 1 and 3 – showed the most remarkable increase in efficacy as compared to the individual drugs.

Here, we carried out a follow-up study utilizing an expanded panel of inhibitors targeting the cell cycle-related kinases as well as several families of proteins responsible for epigenetic modifications. Out of 11 chosen compounds, 6 have been explored in clinical trials and 4 are approved for treatment of solid or haematological cancers (see Supplementary Table S1 for details). We also added a third GB cell line to the study, U-87 MG – the latter is MGMT-negative yet expresses the human apurinic/apyrimidinic endonuclease, which contributes to higher resistance of these cells to ionizing radiation ^13^. The cell lines were incubated with the panel of compounds of interest in normoxic (20% oxygen) or in hypoxic conditions (1% oxygen). Based on the established viability profiles, a PRMT5 inhibitor onametostat was identified as the new player of interest, and its effect was further explored using the spheroid formation assay and transcriptome studies in the onametostat-treated cells (Figure 1).

**Figure 1.**
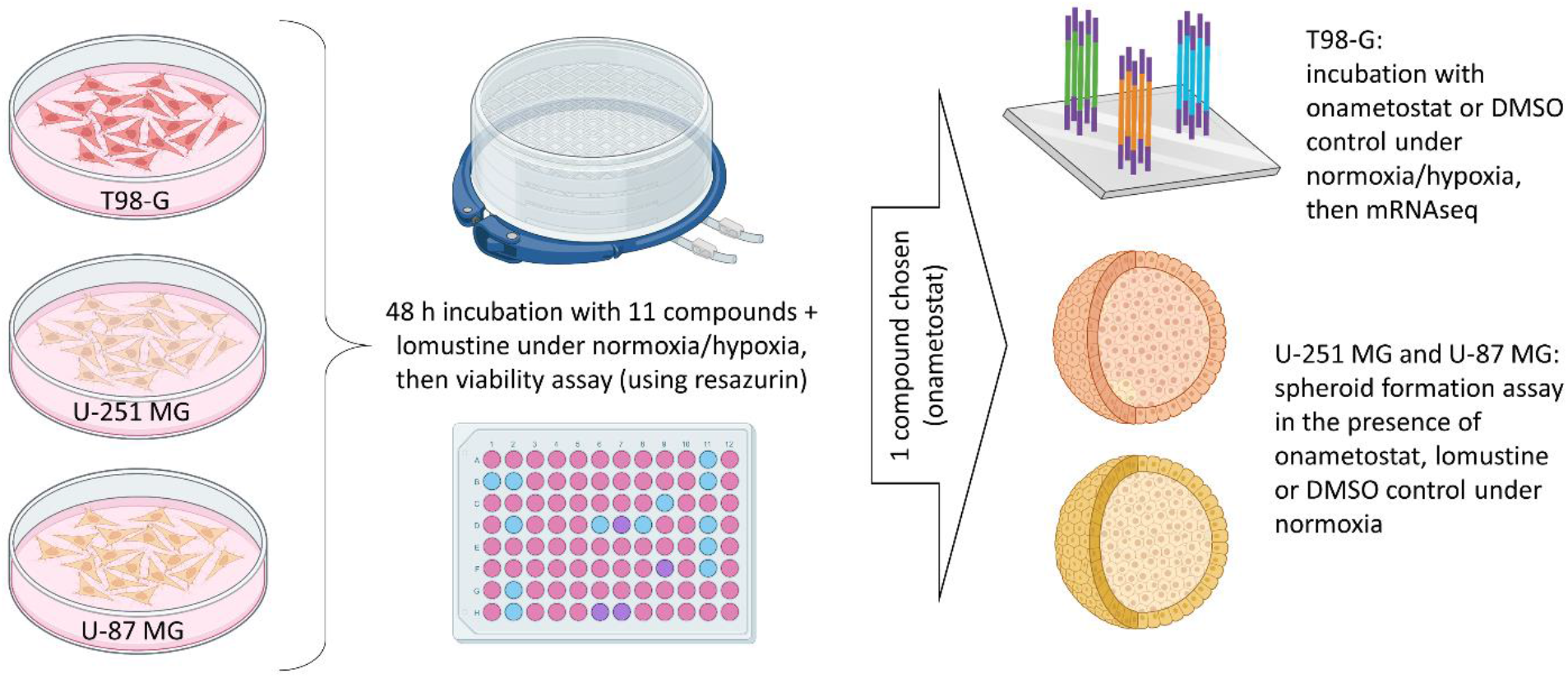
Workflow of the current project. The Figure was composed using BioRender.

## 2. Materials and Methods

### 2.1. Cell lines, chemicals and equipment

Human glioblastoma cell lines T98-G, U-251 MG, and U-87 MG were from the American Type Culture Collection (ATCC; Manassas, VA, USA). Lomustine, AZD1152-HQPA (barasertib), CYC116, danusertib (PHA-739358), MLN8237 (alisertib), MS023, onametostat (JNJ-64619178), palbociclib (PD-0332991) hydrogen chloride salt, tazemetostat (EPZ-6438), and VX 689 (MK-5108) were obtained from Selleckchem (Munich, Germany). Azacytidine and suberanilohydroxamic acid (SAHA, also known as vorinostat) were from Cayman Chemical (Ann Arbor, Michigan, United States). The stock solutions of compounds were prepared in cell culture grade DMSO was from AppliChem (Darmstadt, Germany) and stored at -20 °C.

The solutions and growth medium components for the cell culture were obtained from the following sources: phosphate-buffered saline (PBS), fetal bovine serum (FBS) – Sigma-Aldrich (Steinheim, Germany); Eagle’s Minimum Essential Medium (EMEM) modified to contain Earle’s Balanced Salt Solution, non-essential amino acids, 2 mM L-glutamine, 1 mM sodium pyruvate, and 1500 mg/L sodium bicarbonate – ATCC (Manassas, VA, USA); a mixture of penicillin, streptomycin, and amphotericin B – Capricorn (Ebsdorfergrund, Germany). Resazurin and PBS for viability assay (supplemented with Ca^2+^, Mg^2+^) were from Sigma-Aldrich (St Louis, MO, USA). The propidium iodide for staining of spheroids was obtained from Acros Organics (Switzerland).

The cells were grown at 37 °C in 5% CO_2_ humidified incubator (Sanyo; Osaka, Japan). For viability assay, the initial number of cells was counted using TC-10 cell counter (Bio-Rad; Hercules, CA, USA). In case of the viability assay, the cells were seeded onto transparent 96-well clear flat bottom cell culture plates BioLite 130188; during treatment of cells prior to mRNA sequencing, the cells were grown on the transparent 6-well clear flat bottom cell culture plates BioLite 130184 (both from Thermo Fischer Scientific, Rochester, NY, USA). In case of spheroid formation assay, 96-well black ultra-low attachment spheroid microplates with clear round bottom were used (Corning 4515; Kennebunk, ME, USA).

For hypoxia studies, two different hypoxic chambers were used. In the first case, cultured cells were placed in the modular incubator chamber (Billups-Rothenberg Inc, MIC-101), a flow meter was attached to the unit and the chamber was flushed with 20 L of gas mixture (1% O_2_, 5% CO_2_, 94% N_2_). The chamber was sealed and placed into the incubator (37 °C); the gas mixture was exchanged after every 24 h. In the second case, hypoxic gas mixture (1% O_2_, 5% CO_2_, 94% N_2_) was supplied by gas controller into the incubator of Cytation 5 multi-mode reader system equilibrated at 37 °C (BioTek; Winooski, VT, USA).

Fluorescence intensity and absorbance measurements were carried out with Synergy NEO or Cytation 5 multi-mode readers (both from BioTek; Winooski, VT, USA). Bright-field and fluorescence microscopy was carried out with Cytation 5 multi-mode reader using 4× air objective (1.613 µm/pixel) and automated focussing regime.

### 2.2. Viability assay

T98-G, U-251 MG, or U-87 MG cells (passage number below 15) were seeded in growth medium onto the 96-well plate with the density of 2500, 3500 or 2000 cells per well, respectively (within the linear range of the method, optimized in previous studies ^14^ or shown in Supplementary Figure S1). The cells were left to attach for 24 h at 37 °C in normoxic conditions (95% room air, 5% CO_2_). Next, the growth medium was exchanged, and dilution series of biologically active compounds in PBS were added onto the cells (the final volume of PBS relative to the fetal bovine serum-containing growth medium was 1/10). Based on the solubility of compounds in the water, the following final total concentrations were chosen: azacytidine, lomustine, MS023, SAHA – 6-fold dilution starting from 50 μM; onametostat – 6-fold dilution starting from 20 μM; CYC116, palbociclib, tazemetostat – 6-fold dilution starting from 10 μM; AZD1152-HQPA, danusertib, MLN8237, VX 689 – 6-fold dilution starting from 5 μM. In the case of onametostat and MLN8237, additional dilution series were used for establishing more precise dose-response at low concentrations: starting from 0.3 μM or 200 μM, respectively. An identical volume of PBS was added to the negative control (100% viability). The final volume per well was 200 μL, and the concentration of DMSO in the treated wells was ≤ 0.1% by volume; on each plate, each concentration of each compound was represented in duplicate or triplicate. The cells were incubated for 48 h at 37 °C in normoxic or hypoxic conditions, and viability assay was then carried out according to the previously published protocol ^14^.

### 2.3. Spheroid formation assay

U-251 MG or U-87 MG cells (passage number below 10) were seeded in growth medium onto the 96-well ultra-low attachment plate with the density of 2500 (experiment 1), 2000 (experiment 2) or 1000 (experiment 3) cells per well. At the same time, dilutions of compounds in the growth medium were added to the wells. The final total volume in the well was 200 μL and the final total concentrations were as follows: onametostat – 20 μM, 2 μM, or 0.2 μM; lomustine – 50 μM; DMSO – 0.1% by volume (negative control). The plates were incubated for 48 h at 37 °C in normoxic conditions; next, half of the medium was then replaced with the fresh one (containing the same concentrations of the compounds as outlined above) and the incubation was continued. Following the 95-h incubation, solution of propidium iodide in PBS was added to all wells (final total concentration of 2 mg/mL) and the plates were incubated for 1 h. Finally, imaging was carried out using bright-field (settings – LED intensity 5, integration time 100, detector gain 3) and RFP channel (settings – LED intensity 5, integration time 350, detector gain 15).

### 2.4. Transcriptome studies: sample preparation and mRNA sequencing

T98-G cells (passage number below 10) were seeded in growth medium onto the 6-well plates and grown to 80-85% confluency in normoxic conditions. Subsequently, solutions of onametostat (final total concentration of 1 μM) or DMSO (final total concentration of 0.1% by volume; negative control) in growth medium were prepared and added onto the cells. The treatment lasted for 48 h at 37 °C in normoxic or hypoxic conditions. Next, the medium was removed, the cells were briefly rinsed with PBS and then lysed using RLT buffer (350 μL per well; Qiagen, Hilden, Germany) from the RNeasy Mini kit (Qiagen, Hilden, Germany). The lysates were collected and stored at -80 °C until further use.

The RNA extraction was performed using the RNeasy Mini Kit, together with DNase I treatment (Qiagen, Hilden, Germany), following the manufacturer’s instructions. Afterwards, the RNA concentration of each sample was measured with Nanodrop (IMPLEN, Nordic Biolabs) and Agilent TapeStation QC (Agilent Technologies Inc., CA, USA). High-quality samples (A260/A280 > 1.8, and RNA integrity > 9) were selected for library preparation using the Illumina Stranded mRNA Prep Ligation protocol (Illumina, USA), following the manufacturer’s instructions, and using 200 ng of RNA as input. The final library included 12 samples, and it was sequenced using the Illumina NextSeq2000 sequencing platform 100 cycle flow cell in Bioinformatics and Expression Analysis core facility (BEA) at Karolinska Institute, Sweden. For sequencing, 1000 pM of the library was loaded on the flow cell.

### 2.5. Statistical analysis and software

For general data analysis, GraphPad Prism 6 (San Diego, CA, USA) and Excel 2016 (Microsoft Office 365; Redmond, WA, USA) were used. Throughout the study, the grouped comparisons were carried out using one-way ANOVA with Dunnett’s test for multiple comparisons; unless indicated otherwise, the pairwise comparisons were carried out using the unpaired two-tailed t-test with Welch’s correction. In all statistical tests, the significance of comparisons is indicated as follows: *** indicates P ≤ 0.001, ** indicates P ≤ 0.01,* indicates P ≤ 0.05.

In case of viability assay, the data analysis was carried out according to the previously published protocol ^14^. Specifically, in each independent experiment, the fluorescence intensity measured for the replicate treatments was pooled and the data obtained for the negative control was plotted against incubation time with resazurin. One time-point within duration of data acquisition was chosen where the signal of the negative control remained in the linear range, and only data measured at this time-point was used for the further analysis. For normalization, data obtained for wells treated with PBS (blank control) was considered as 100% viability; data acquired for the 50 μM resazurin solution (in the absence of cells) was considered as 0% viability. Next, the ratio of absorbance at 570 μm and 600 μm was calculated for each well. The ratios were analyzed analogously to the fluorescence intensity data, and the normalized viability values calculated from the fluorescence intensity and the absorbance measurements were pooled. Finally, data from all independent experiments (N ≥ 3 for each treatment in each cell line) was pooled for each individual compound and oxygenation condition. The pooled normalized viability was plotted against the concentration of compound in the dilution series and fitted to the logarithmic dose-response function (three parameters), or to the biphasic equation with the Hill slope values fixed at -1 and the fraction of the curve derived from the more potent phase fixed between 0 and 1.

In spheroid formation assay, ImageJ software (Fiji package ^15^) was used for the image analysis. The spheroid contour was denoted manually using the freehand selections tool and the area, diameter and circularity of the spheroid were quantified; in case of disintegrated spheroids with poorly defined borders, the selection involved all dark area covered with cells. For the RFP channel images, the average signal intensity of propidium iodide per unit of area was also established. The measured parameters were normalized to the negative control in the given cell line (set to 100%) in each independent experiment (N = 2 for each treatment), and the data for the identically treated cells within the same cell line was then pooled for all independent experiments.

The RNA sequencing reads were initially assessed for quality using the FASTQC program (version 0.11.8) to ensure high quality data ^16^. Based on the FASTQC results, MultiQC program was utilized to generate a comprehensive report on the sequencing data quality ^17^. Subsequently, the reads underwent trimming using the Fastp program ^18^. The trimmed reads were then aligned to the human reference genome GRCh38 using STAR 2.7.5a ^19^. To estimate the read counts per gene, we utilized the featureCounts tool with default parameters ^20^.

We implemented the T-Distributed Stochastic Neighbor Embedding (t-SNE) technique for clustering analysis using the R package Rtsne ^21^. Prior to t-SNE, we applied the variance stabilizing transformation to the raw count data using the DeSeq2 Bioconductor package ^22^. This transformation was carried out to eliminate potential biases and enhance the accuracy of the clustering analysis.

Differential gene expression analysis was performed in the R environment using the edgeR 3.28 package ^23^. Genes with low counts were filtered out prior to the differential gene expression analysis using the filterByExpr function from the edgeR package to ensure robust results. Genes with absolute log fold change |logFC| > 1 and FDR-adjusted P-value (q-value) < 0.05 were considered as differentially expressed.

To create the Venn diagram and the Volcano plots, we utilized the interactive tool Venny ^24^ and the EnhancedVolcano R package ^21^, respectively. For analysis of the cellular pathways up- or downregulated in various treatment comparisons, an online platform Metascape was applied (version V3.5.20230501 ^25^).

## 3. Results

### 3.1. Viability profiling of GB cell lines following treatment in normoxic or hypoxic conditions

As the first step of our study, we established the viability profiles of T98-G, U-251 MG and U-87 MG cell lines following the 48-h incubation with the set of twelve compounds under normoxic or hypoxic conditions. The compounds included 5 inhibitors of enzymes carrying out epigenetic modifications (including DNMT family, histone-lysine N-methyltransferase EZH2, protein arginine methyltransferase family PRMT, histone deacetylase family HDAC), 6 inhibitors of the cell cycle-related protein kinases (including Aurora and cyclin-dependent kinase CDK families), and a control compound lomustine. The details on the previously reported targets of the compounds of interest are listed in the Supplementary Table S1; apart from MS023, all compounds have been tested in clinical trials or approved for clinical use.

The negative logarithms of IC_50_ values (pIC_50_) of compounds measured in the viability assay are summarized in Table 1, and visualized as radar plots in Figure 2 and Supplementary Figure S2. The dose-response curves of individual compounds are shown in Supplementary Figures S3-S5; additional parameters (apart from the IC_50_ values) obtained from the dose-response curves are compared in Supplementary Figure S6. Overall, T98-G was found to be the least and U-87 MG the most chemosensitive cell line, yet the same compounds were identified as the most efficient in all cell lines tested: an Aurora A kinase inhibitor MLN8237, and a PRMT5 inhibitor onametostat. Both compounds featured biphasic dose-response profile, with the low-dose IC_50_ values in low nanomolar to subnanomolar range. In accordance with the previous study ^12^, an Aurora A/B inhibitor AZD-1152 HQPA was also found highly efficient in U-251 MG cells.

**Table 1.**
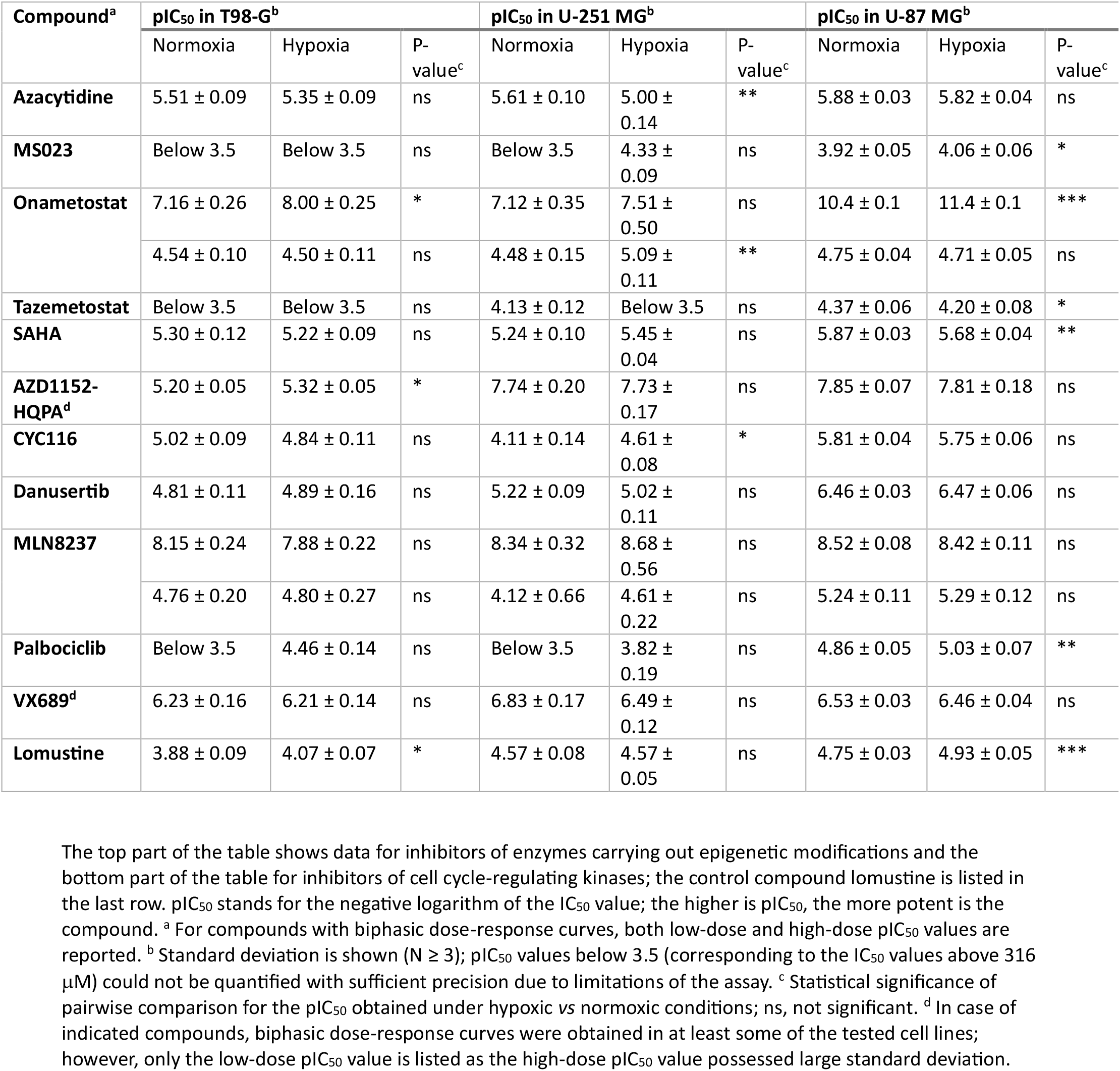
Viability assay data following incubation of GB cell lines with compounds for 48 h under normoxic or hypoxic conditions.

**Figure 2.**
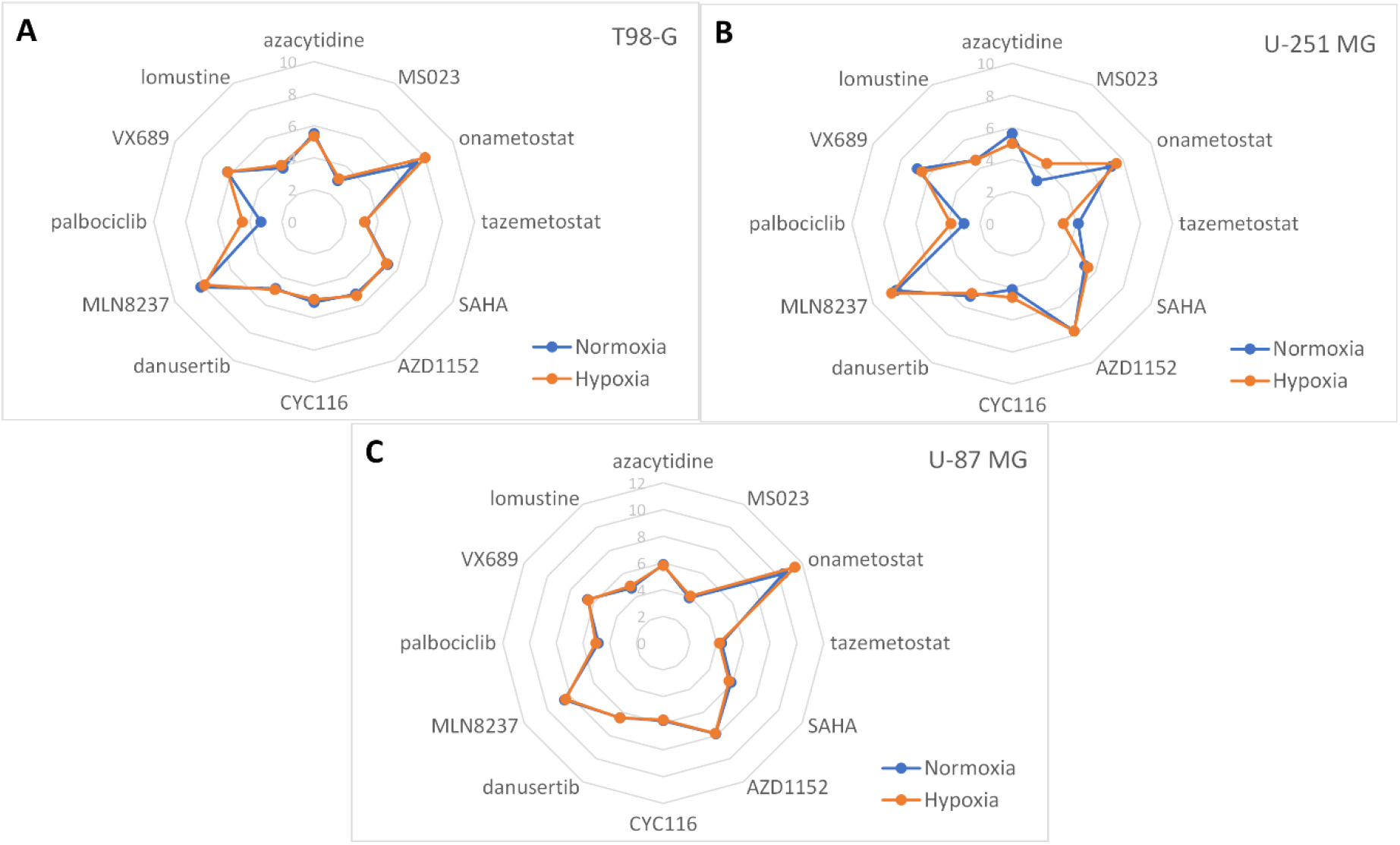
Radar plots enabling comparison of viability pIC_50_ values obtained for different compounds following incubation of cells under normoxic or hypoxic conditions. Cell lines: (A) T98-G, (B) U-251 MG, (C) U-87 MG; the incubation conditions are shown in the right bottom corner. The names of compounds are listed along the radar perimeter. The data was obtained by pooling all independent experiments (N ≥ 3). For clarity, no error bars are depicted and only one pIC_50_ value is shown per compound (in case of compounds featuring the biphasic dose-response fit, only the largest pIC_50_ value was chosen). The numbering of y-axis is shown in light grey. Please note that the y-scale ranges from 0 to 10 in panels A-B, and from 0 to 12 in panel C.

The effect of hypoxia was relatively small, yet statistically significant differences in compound efficacy under hypoxic *vs* normoxic conditions were observed in several cases. According to the comparison of IC_50_ values, under hypoxic conditions the efficacy of 5-azacytidine was reduced in U-251 MG cells (P < 0.01) and the efficacy of SAHA in U-87 MG cells (P < 0.01). The efficacy of tazemetostat was reduced in U-87 MG (P < 0.05) as well as in U-251 MG cells, but in the latter case, the exact IC_50_ value under hypoxic conditions was too high to measure with the assay used. On the other hand, increase of efficacy under hypoxic conditions could be observed for onametostat in all cell lines (P < 0.05), for AZD1152-HQPA in T-98G cells (P < 0.05), for CYC116 in U-251 MG cells (P < 0.05), and for lomustine in both T-98G and U-87 MG cells (P < 0.05 and P < 0.001, respectively). In case of MS023 and palbociclib, the trend toward efficacy increase under hypoxia was evident in all cell lines, yet due to the generally high IC_50_ values of these compounds, statistical significance could only be confirmed in U-87 MG cells (P < 0.05 for MS023 and P < 0.01 for palbociclib).

### 3.2. Onametostat affects formation of U251 and U87MG spheroids

As onametostat showed the most promising trends from the aspect of both low IC_50_ and efficacy increase in hypoxia, we proceeded with more detailed characterization of this compound. In U-251 MG and U-87 MG cell lines, we explored the effect of onametostat on the formation of tumour spheroids. Increasing concentrations of onametostat (final total concentration of 0.2, 2, or 20 μM) or control compounds (positive control: 50 μM lomustine, negative control: 0.1% DMSO) were added onto the ultra-low attachment microplates together with the cells. The concentrations of onametostat were chosen taking into consideration the trends observed in the viability assay, sufficiently low to avoid the excessive cell death yet sufficiently high to make the effect observable even in 3D culture, which is usually less sensitive to compound treatment than 2D ^26–28^. The size of spheroids and the intensity of propidium iodide (PI) staining in necrotic/late apoptotic cells with compromised membrane integrity was assessed after 96 h of incubation. As the inner part of the tumour spheroids permanently resides at the hypoxic conditions due to the limited diffusion of oxygen ^29^, only normoxic incubation conditions were used.

The results of the assay are summarized in Figure 3 and Supplementary Figure S7. In general, the interrupting effect of a compound on the spheroid formation can manifest itself either as decrease of spheroid size (if the compound affects viability and/or proliferation of the cells) or apparent increase of spheroid size (if the compound weakens the cell-cell contacts). Indeed, both trends were evident in our data. In U-251 MG cells, both onametostat and lomustine caused significant increase of the spheroid diameter (P < 0.05), with slight dose-response effect observable in case of onametostat (*i.e*., more concentrated compound resulting in larger apparent diameter). On the other hand, in U-87 MG cells, reduction of spheroid diameter was evident in case of highest onametostat concentration, while lowest concentration of the compound caused apparent increase in spheroid size (P < 0.001). Such difference in the cell line behaviour could be explained by the higher proliferation rate but also higher chemosensitivity of the U-87 MG cells, which was also previously observed in the viability assay with the adherent cells (Table 1). Furthermore, increase of the PI staining intensity (normalized by the spheroid area) was observable in U-87 MG spheroids treated with 20 μM onametostat.

**Figure 3.**
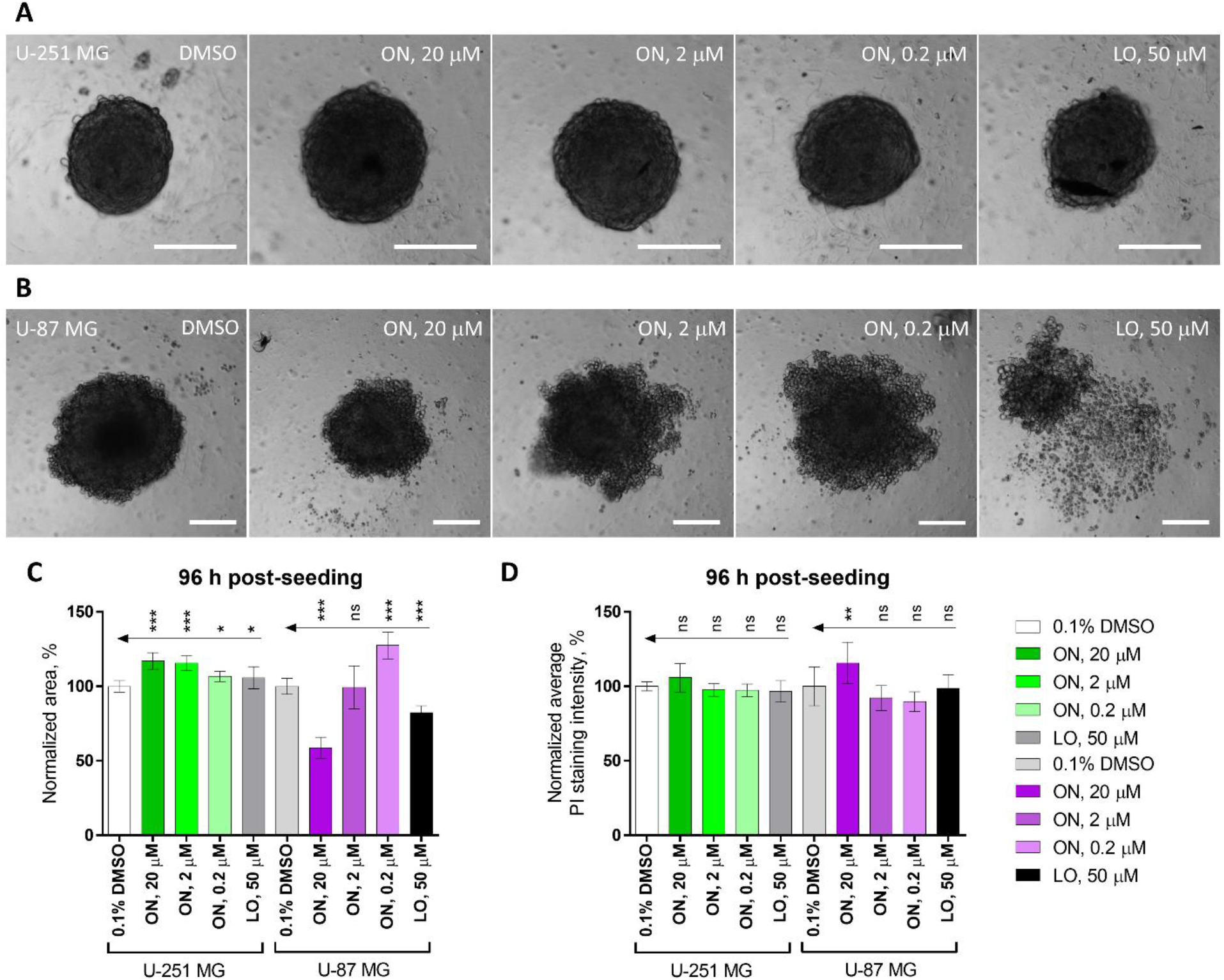
Effect of onametostat (ON) or lomustine (LO) on the spheroid formation in glioblastoma cell lines. Panels (A) and (B) show examples of spheroid morphology in U-251 MG cells and U-87 MG cells, respectively, following the 96-h treatment with indicated compounds in a single representative experiment; scale bar: 200 μM. Panel (C) summarizes normalized spheroid area and panel (D) summarizes normalized average intensity of propidium iodide staining per spheroid area using data pooled from 2 independent experiments (a total of 11-12 spheroids per condition); error bars indicate standard deviation. The grouped comparisons show statistical significance of differences for the parameters measured following incubation of cells with onametostat or lomustine *versus* 0.1% DMSO (one-way ANOVA): *** indicates P ≤ 0.001, ** indicates P ≤ 0.01, * indicates P ≤ 0.05, ns indicates not significant.

### 3.3. Transcriptome studies in T98-G lysates: establishing the characteristic hypoxia and onametostat signatures

As the final part of our experimental effort, we explored the effect of 1 μM onametostat on the transcriptome in T98-G cell line. The adherent cells were incubated with the compound for 48 h under normoxic or hypoxic conditions; as a negative control, 0.1% DMSO was again used. T98-G was chosen as it was overall the least chemosensitive among the cell lines screened in this study (Figure 2).

For different samples within 3 independent experiments, a read count distribution ranging from 16.8 million to 58.2 million reads was observed. The majority of samples exhibited read counts within the interquartile range (IQR) of 23.3 million to 53.9 million reads, indicating a consistent level of coverage across the majority of the samples (Supplementary Table S4). The t-distributed stochastic neighbor embedding plot (t-SNE; Figure 4) illustrates the clustering based on the similarity of the transcriptome profiles obtained for the different treatments. It is evident that the impact of onametostat treatment is markedly more pronounced than the effect of oxygenation conditions; still, the clustering is distinguishable between the hypoxic vs normoxic conditions, and the independent experiments cluster together.

**Figure 4.**
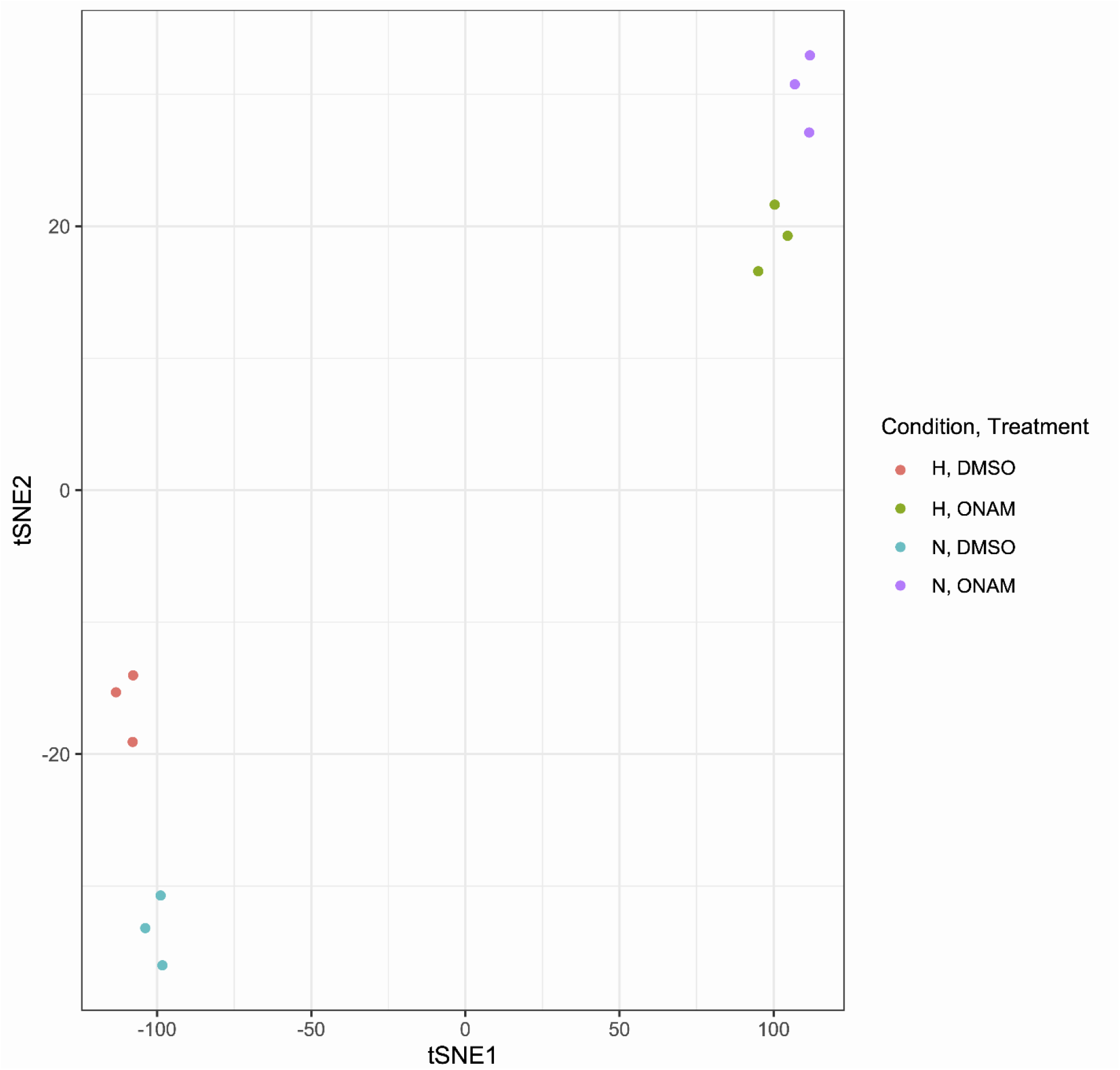
Clustering shows differentiation between the treatments (N = 3). The t-SNE plot is based on the entire transcriptome data after filtering out genes with low counts. The colour codes for different oxygenation conditions and treatments are shown on the right. Abbreviations: H, hypoxia; N, normoxia; ONAM, onametostat.

Figure 5A lists the number of the differentially expressed genes (DEG) in pairwise comparisons of different treatments; FDR rate of 0.05 was used as a cut-off for defining the statistically significant differences. The overall number of both up- and downregulated DEGs is lower in comparisons of cells treated under hypoxic *vs* normoxic conditions than the number of DEGs in comparisons of cells treated with onametostat *vs* DMSO. This is consistent with the more pronounced effect of PRMT5 inhibition as compared to the oxygen deprivation. The number of DEGs common for the various comparisons are shown in the Venn diagram in Figure 5B. In line with the t-SNE plot, the cells treated with onametostat share a number of DEGs relative to the DMSO-treated cells irrespective of the oxygen content during the treatment. The lists of DEGs termed significant for the different treatment comparisons (FDR < 0.05) are provided in the Supplementary Table S2A-H.

**Figure 5.**
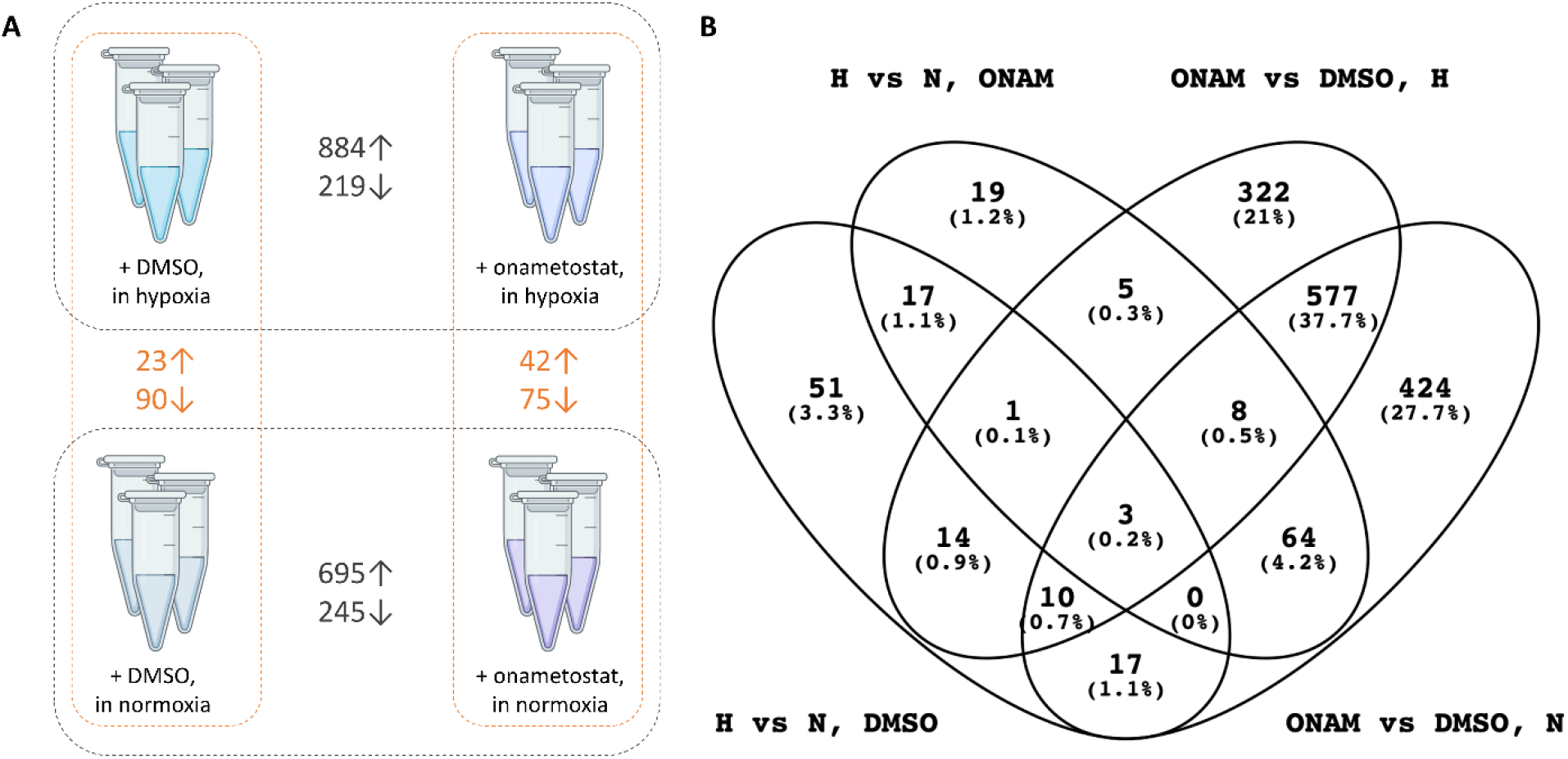
DEG counts in treatment comparisons and number of common DEGs in various treatments. (A) Number of DEGs identified as significantly enriched in the pairwise comparisons of differently treated cells (FDR < 0.05); ↑ indicates higher abundance in hypoxia- or onametostat-treated cells and ↓ indicates higher abundance in normoxia- or DMSO-treated cells. (B) Venn diagram of the DEGs in various treatment comparisons (FDR < 0.05 in any compared treatments). Abbreviations: H, hypoxia; N, normoxia; ONAM, onametostat.

We next focused specifically on the two types of comparisons: first, DEGs observed for the hypoxic *vs* normoxic conditions in the DMSO-treated cells (Volcano plots shown in Figure 6A), and second, DEGs observed for the onametostat *vs* DMSO treatment in hypoxia (Volcano plots shown in Figure 6B). The cellular pathways to which the established DEGs belong were analysed using Metascape, an online platform that integrates data from the following ontology sources: KEGG Pathway, GO Biological Processes, Reactome Gene Sets, Canonical Pathways, CORUM, and WikiPathways ^25^. The major results are summarized in Table 2 and shown in detail under the Supplementary Table S3A-D. Additionally, for visualization of differences of onametostat effect under different oxygenation conditions, the Volcano plot featuring DEGs observed for the onametostat *vs* DMSO treatment in normoxia is provided in the Supplementary Figure S8.

**Figure 6.**
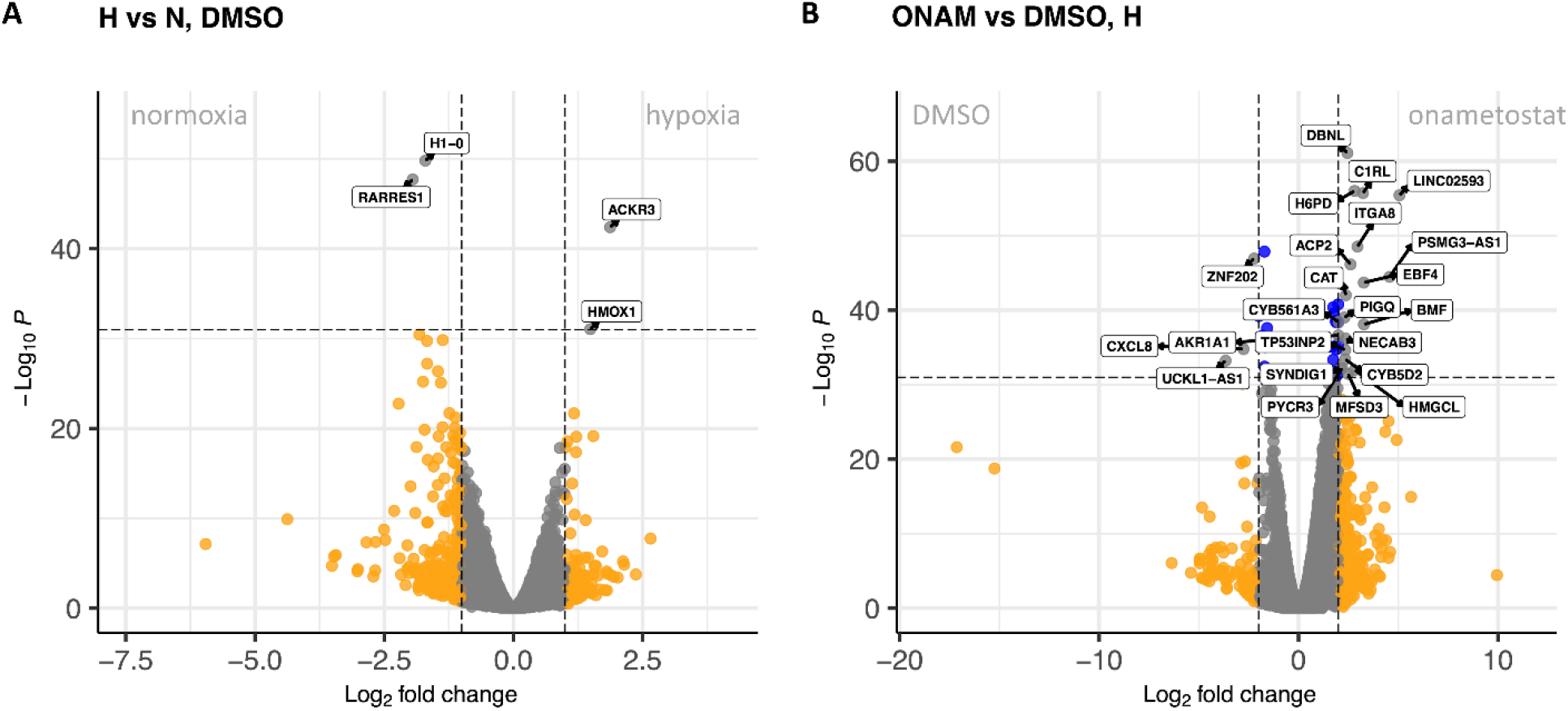
Volcano plots showing DEGs in treatment comparisons. (A) Hypoxic *vs* normoxic conditions in the T98-G cells treated with 0.1% DMSO; top hits are marked with the name labels, and DEGs coloured in orange possessed binary logarithm of fold change values of below -1 or over 1. (B) Onametostat-*vs* DMSO-treated T98-G cells following incubation in hypoxia. Top hits are marked with the name labels; DEGs coloured in orange possessed binary logarithm of fold change values of below -2 or over 2, while DEGs coloured in blue featured the P-value cut-off of 10^−32^. Abbreviations: H, hypoxia; N, normoxia; ONAM, onametostat.

**Table 2.**
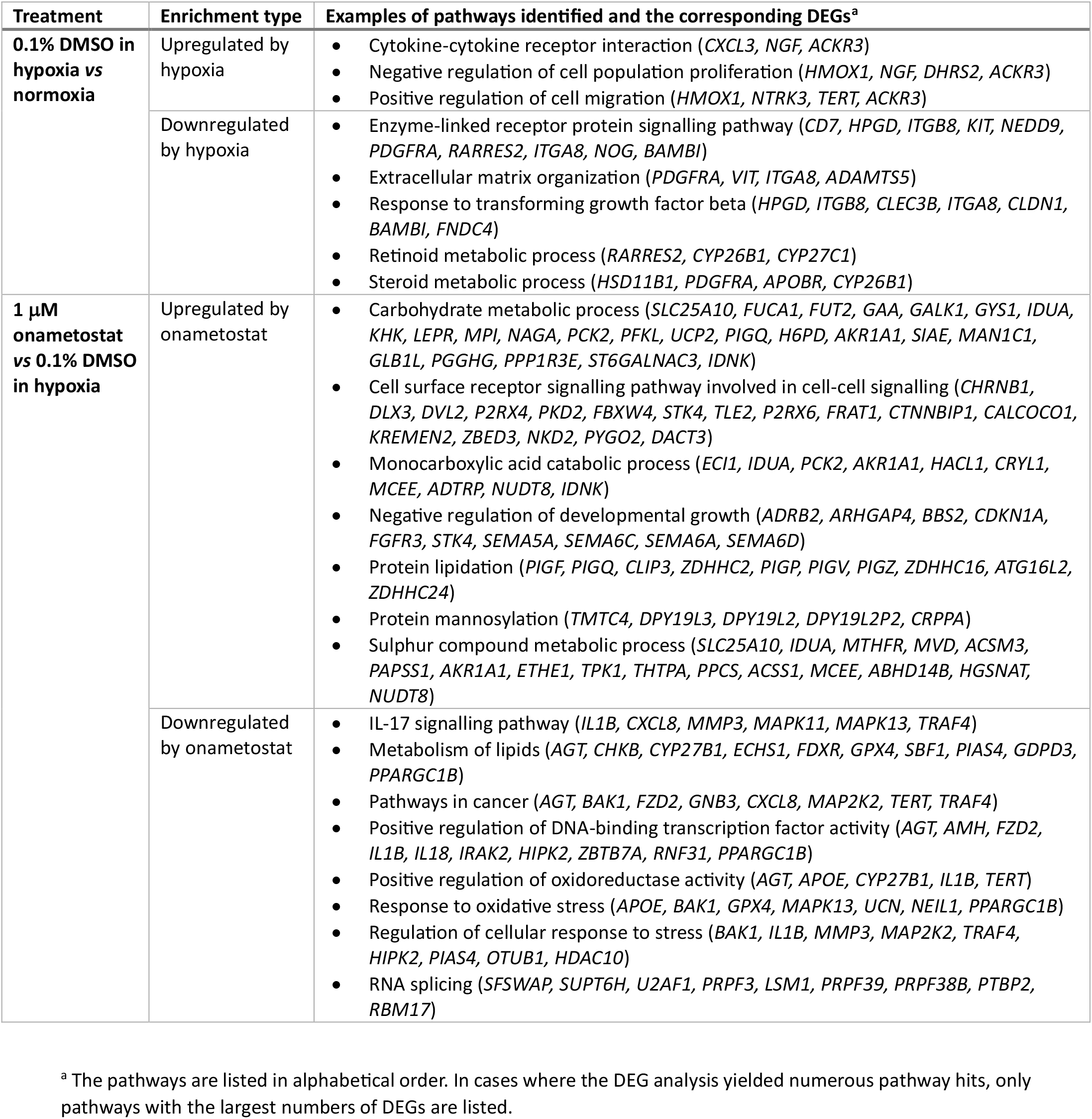
Cellular pathways identified by the Metascape platform based on the DEGs corresponding to the comparisons of the differently treated samples (FDR < 0.05).

Overall, in T98-G cells, 48-h incubation under hypoxic conditions triggered reorganization of the extracellular matrix (ECM), enhanced cell migratory properties, and altered several metabolic processes. Treatment with onametostat also resulted in metabolic changes, yet it also reduced activity of various transcription factors, suppressed RNA splicing, and caused overall downregulation of the signalling pathways sustaining survival of the cancer cells.

## 4. Discussion

The current study served as a follow-up to the previously published effort where the viability profile of GB cell lines U-251 MG (MGMT-negative) and T98-G (MGMT-positive) was established following inhibition of various pathways putatively crucial for the survival of the cancer cells ^12^. Here, we expanded the choice of inhibitors targeting the two types of hits identified previously: the cell cycle-regulating kinases and the epigenetic writers or erasers. To provide deeper insight into the variations in compound efficacy dependent on the tumour- or environment-dictated factors, we additionally included a third cell line and explored the effect of short-term hypoxia treatment (48 h).

The viability assay indicated relatively small effect of hypoxia on the efficacy of compounds in all tested cell lines, with the largest shift of dose-response curves observed for MS023 in U-251 MG cells, palbociclib in T98-G cells, and onametostat in T98-G cells (all potentiated by hypoxia; see Supplementary Figures S3-S5). The moderate effect of hypoxic treatment on the gene transcription in the DMSO-treated T98-G cells confirmed this notion (Figure 4). In literature, a recent study in the same cell line reported a substantial number of DEGs following the 72-h hypoxia treatment of passage number 3 cells, with upregulation of the Warburg effect-related genes such as inositol-requiring enzyme 1 (*IRE1*) or lactate dehydrogenase A (*LDHA*) ^30^. Neither IRE1 nor LDHA were identified as DEGs in our study, which may be explained by the shorter treatment times and higher cell passage number in our experiments. The longer culturing of cells was chosen due to our previous observation that in case of lower passages (below number 4), the proliferation rate of the cells changes remarkably (mirrored by the apparent drop in efficacy of the Aurora A inhibitors) ^14^, and we thus chose later passages (number 8-9) for the transcriptomic study to minimize the putative effect of the proliferation rate change.

Still, several DEGs short-listed in our study (Table 2 and Supplementary Table S2) have been previously linked to hypoxia. Among the genes upregulated in hypoxia in both DMSO- and onametostat-treated cells, the atypical chemokine receptor 3 (*ACKR3*) has been explored in the context of cardiovascular diseases and inflammation ^31^, the nerve growth factor (*NGF*) in the context of angiogenesis in non-small cell lung cancer ^32^, and the 4-hydroxyphenylpyruvate dioxygenase (*HPD*) in the context of metabolic reprogramming in lung cancer ^33^. In case of the hypoxia-upregulated DEGs found only in the DMSO-treated cells, the heme oxygenase-1 (*HMOX1*) and isthmin1 (*ISM1*) have been reported as the direct downstream targets of the hypoxia-inducible transcription factor (HIF) ^34,35^, whereas indirect regulation mechanism by HIF has been proposed for the C-X-C motif chemokine ligand 3 (*CXCL3*) ^36^. From the hypoxia-induced pathways identified by the Metascape algorithm (Table 2), upregulation of the cytokine stimulation and increase of cell migratory properties as well as remodelling of the ECM can be directly linked to inflammation and metastasis formation in cancer ^37–39^. Somewhat surprisingly, Metascape also identified the cluster corresponding to the response to transforming growth factor β (TGF-β) among the pathways downregulated under hypoxic conditions in our study. In general, HIF is known to activate the TGF-β signalling ^40^ – thus, our current finding might point to reorganization rather than suppression of the pathway under hypoxic conditions.

Among the tested compounds that affected the cell viability at two-digit nanomolar or lower concentration in both normoxia and hypoxia across the cell lines (Table 1), we chose PRMT5 inhibitor onametostat as the compound of interest for the further studies. According to literature, inhibition of PRMT5 has been termed an efficient strategy in cancers featuring deletion of the methylthioadenosine phosphorylase gene (*MTAP*), as such cancers rely on PRMT5 for survival ^41^. Importantly, in GB, *MTAP* is one of the most frequently deleted genes ^42^ – although it has been shown that the success of PRMT5 inhibition in the context of GB is also dependent on other factors ^43^.

While we are not aware of previous reports on the effect of onametostat on the GB cell line transcriptome, the list of DEGs in onametostat-*vs* DMSO-treated cells revealed several players previously reported as characteristic of PRMT5 inhibition (Table 2 and Supplementary Table S2). For instance, PRMT5 is known to regulate DNA repair and mRNA splicing ^44^, both of which were short-listed as downregulated pathways in our data by the Metascape algorithm. Among the DEGs belonging to the mRNA splicing pathway, polypyrimidine tract-binding protein 2 (*PTBP2*) and U2 small nuclear RNA auxiliary factor 1 (*U2AF1*) have been previously shown to promote proliferation in glioma ^45,46^. The altered expression of several other genes following the treatment of T98-G cells with onametostat in both normoxic and hypoxic conditions also indicates interference of the compound with the cancer survival strategies. Among the downregulated DEGs, the pleckstrin homology domain containing S1 (*PLEKHS1*) and the cluster representing the IL-17-regulated genes are of interest as these can be linked to the PI3K/Akt pathway ^47,48^, which is frequently mutated in various solid cancers, including GB ^49^. Additionally, the semaphorins 5A and 6A (*SEMA5A, SEMA6A*) which we identified among the upregulated DEGs have been shown to act as the potential suppressors of cancer migration ^50,51^.

Furthermore, several onametostat treatment-related DEGs indicate the cell cycle arrest at G1 phase: e.g., the histone cluster downregulated in both normoxic and hypoxic conditions ^52^, the early B-cell transcription factor 4 (*EBF4*) upregulated in both normoxic and hypoxic conditions ^53^, and the cell cycle inhibitor p21 (*CDKN1A*) upregulated in hypoxic conditions ^54^. The cell cycle arrest is in line with the biphasic dose-response curves of onametostat observed in viability assay for all cell lines (Supplementary Figures S3C, S4C, S5C), as the low-dose effect is frequently indicative of cytostatic rather than cytotoxic effects ^14^.

Finally, several of the onametostat-upregulated DEGs highlight the survival strategies of GB and cancer in general, as the post-treatment transcriptome inevitably includes the gene pool from the surviving population of the T98-G cells. Such DEGs, found upregulated in both normoxic and hypoxic conditions, include hexose-6-phosphate dehydrogenase (*H6PD*) that contributes to metabolic reprogramming in glioma stem cells ^55^, protein tyrosine kinase 2b (PTK2B) that enhances glioma cell migration ^56^, and catalase (*CAT*) that regulates chemo- and radioresistance in glioblastoma ^57^. Our short-list of DEGs did not contain the microtubule regulator stathmin 2 that was reported as the PRMT5 inhibitor resistance-promoting player in a different cancer context (lung adenocarcinoma) ^58^. However, we found that another microtubule-related molecular target was upregulated by onametostat under hypoxic conditions (microtubule-associated serine/threonine kinase 1, *MAST1*), which has also previously been linked to the putative resistance mechanisms in cancer ^59^.

The given study was limited to the *in vitro* settings, and our major findings – whether related to sensitivity or resistance mechanisms – should thus be confirmed in clinically more relevant models. We have also not addressed the blood-brain barrier (BBB)-penetrative properties of the tested compounds. However, it has been shown that the integrity of BBB is disrupted following radiotherapy ^60,61^, which can be utilized for development of combined treatment schemes. As onametostat induces cell cycle arrest at the G1 phase, which represents the less sensitive cell cycle phase to radiotherapy ^62,63^, it is not advisable to apply it prior to radiation, but rather following the latter. Furthermore, despite the emergence of DEGs indicating resistance to onametostat in our transcriptome study, it is worth to investigate effect of mixtures of onametostat with lomustine for achieving improved effect on proliferation of GB cells at low doses of the PRMT5 inhibitor. Overall, we confirmed that inhibition of PRMT5 is a promising strategy that should be explored more intensely in the context of treatment of recurrent glioblastoma.

## Acknowledgements

We thank Ago Rinken group members for the maintenance of the microplate reader and microscopy platform.

## Author Contributions

Conceptualization, D.L. and J.J.; Methodology, D.L., M.K.K., K.-L.E. and V.M.; Validation, D.L., M.K.K., H.V.,

V.D.N. and H.L.; Formal Analysis, D.L. and M.K.K.; Resources, D.L., V.D.N., A.S. and J.J.; Writing – Original Draft Preparation, D.L., V.M. and V.D.N.; Writing – Review & Editing, D.L., M.K.K., H.V., V.M., V.D.N., H.L., K.-L.E., A.S. and J.J.; Visualization, D.L. and V.M.; Supervision, D.L., K.-L.E., A.S. and J.J.; Project Administration, A.S. and J.J.; Funding Acquisition, A.S. and J.J.

## Data availability

The mRNA sequencing raw data are available in Gene Expression Omnibus (GEO) at https://www.ncbi.nlm.nih.gov/geo/ and can be accessed with GEO accession number: GSE235648. Other datasets generated and analysed during the current study are available from the corresponding author on reasonable request.

