## Supplementary for "Inhibition of epigenetic and cell cycle-related targets in glioblastoma cell lines: onametostat reduces proliferation and viability in both normoxic and hypoxic conditions"

**SUPPLEMENTARY DATA**

**Supplementary methods**

***Dynamic range optimization of resazurin assay with U-87 MG cells***

Two-fold dilution series of cell suspension in the growth medium (containing FBS) were carried out directly on the 96-well flat-bottom plates. The cells were first grown for 24 h in the $CO_2$ incubator at 37 °C under normoxic conditions and subsequently for further 48 h in the $CO_2$ incubator (37 °C) under normoxic or hypoxic conditions. The total volume of liquid per well was 200 μL. Afterwards, the growth medium was removed, the cells were rinsed with PBS, and solution of 50 μM resazurin in PBS (supplemented with $Ca^{2+}$ and $Mg^{2+}$) was added. The plate was immediately transferred to the microplate reader (Biotek NEO or Cytation 5) and readings were taken at 37 °C in the kinetic mode (every 15 min for 90-105 min) using the following parameters: (A) fluorescence intensity: excitation 540 nm, emission 590 nm, mono-chromator, top optics, gain 50, slit width 15 nm in case of NEO and 20 nm in case of Cytation 5; (B) absorbance at 570 nm and 600 nm, monochromator; read height 8.5 mm. A total of 3 independent experiments were performed.

Table S1. Information on the targeted compounds applied in the study, their biological targets and clinical use

| Compound | Target protein | Cellular processes affected | Status in clinics[a] |
|---|---|---|---|
| **Azacytidine** | DNA methyltransferase 1 and 3 (DNMT1, DNMT3) | Epigenetic modifications | Approved for treatment of myelodysplastic syndrome in the USA and EU and of the juvenile myelomonocytic leukaemia in the USA |
| **MS023** | Type 1 protein arginine methyltransferases (PRMT) | Epigenetic modifications | Pre-clinical, available from the Structural Genomics Consortium's (SGC) Epigenetics Probes Collection |
| **Onametostat** | Protein arginine methyltransferase 5 (PRMT5) | Epigenetic modifications | Clinical trials, Phase 1 (relapsed/refractory B cell non-Hodgkin lymphoma or advanced solid tumours) |
| **Tazemetostat** | histone-lysine N-methyltransferase enhancer of zeste homolog 2 (EZH2) | Epigenetic modifications | Approved in the USA for treatment of metastatic or locally advanced epithelioid sarcoma and of the relapsed or refractory follicular lymphoma |
| **SAHA** | Histone deacetylase 1, 2 and 3 (HDAC1, HDAC2, HDAC3) | Epigenetic modifications | Approved in the USA for treatment of progressive, persistent, or recurrent cutaneous T cell lymphoma |
| **AZD1152-HQPA** | Protein kinase Aurora B (AurB) | Mitosis | Clinical trials, Phase 1 (relapsed/refractory acute myeloid leukaemia) |
| **CYC116** | Protein kinases Aurora A, B (AurA, AurB) and vascular endothelial growth factor receptor 2 (VEGFR2) | Mitosis | Clinical trials, Phase 1 (advanced solid tumours) |
| **Danusertib** | Protein kinases Aurora A, B and C (AurA, AurB, AurC) | Mitosis | Clinical trials, Phase 2 (hormone refractory prostate cancer; multiple myeloma; and relapsed chronic myelogenous leukemia) |
| **MLN8237** | Protein kinase Aurora A (AurA) | Mitosis | Clinical trials, Phase 2 (relapsed/refractory peripheral T-cell lymphoma; and unresectable stage III-IV melanoma) |
| **Palbociclib** | Protein kinases cyclin-dependent kinases 4 and 6 (CDK4, CDK6) | Mitosis | Approved in the USA for treatment of advanced (metastatic) breast cancer |
| **VX 689** | Protein kinase Aurora A (AurA) | Mitosis | Clinical trials, Phase 1 (advanced and/or refractory solid tumours) |

[a] Sources of data: U.S. National Library of Medicine https://clinicaltrials.gov/ct2/home (last accessed May 8, 2023) and IUPHAR/BPS https://www.guidetopharmacology.org/ (last accessed May 8, 2023)

Table S2. Differentially expressed genes (DEGs; FDR < 0.05) in various treatment comparisons

Provided as a separate Excel file:

- Part A: DEGs upregulated in hypoxic *vs* normoxic conditions, cells treated with 0.1% DMSO
- Part B: DEGs downregulated in hypoxic *vs* normoxic conditions, cells treated with 0.1% DMSO
- Part C: DEGs upregulated in hypoxic *vs* normoxic conditions, cells treated with onametostat
- Part D: DEGs downregulated in hypoxic *vs* normoxic conditions, cells treated with onametostat
- Part E: DEGs upregulated in onametostat- *vs* DMSO-treated cells under normoxic conditions
- Part F: DEGs downregulated in onametostat- *vs* DMSO-treated cells under normoxic conditions
- Part G: DEGs upregulated in onametostat- *vs* DMSO-treated cells under hypoxic conditions
- Part H: DEGs downregulated in onametostat- *vs* DMSO-treated cells under hypoxic conditions

Abbreviations: logFC, binary logarithm of fold change; logCPM, logarithm of counts per million; FDR, false discovery rate.

Table S3. Full lists of the cellular pathways identified by the Metascape platform based on the DEGs corresponding to the comparisons of the differently treated samples (FDR < 0.05)

Provided as a separate Excel file:

- Part A: pathways upregulated in hypoxic *vs* normoxic conditions, cells treated with 0.1% DMSO
- Part B: pathways downregulated in hypoxic *vs* normoxic conditions, cells treated with 0.1% DMSO
- Part C: pathways upregulated in onametostat- *vs* DMSO-treated cells under hypoxic conditions
- Part D: pathways downregulated in onametostat- *vs* DMSO-treated cells under hypoxic conditions

All genes in the genome have been used as the enrichment background. Terms with a p-value < 0.01, a minimum count of 3, and an enrichment factor > 1.5 (the enrichment factor is the ratio between the observed counts and the counts expected by chance) are collected and grouped into clusters based on their membership similarities. Count is the number of genes in the user-provided lists with membership in the given ontology term. % is the percentage of all user-provided genes that are found in the given ontology term (only input genes with at least one ontology term annotation are included in the calculation). Log10(P) is the p-value in log base 10. Log10(q) is the multi-test adjusted p-value in log base 10. The p-values are calculated based on the cumulative hypergeometric distribution, and the q-values are calculated using the Benjamini-Hochberg procedure to account for multiple testings.

Table S4. mRNA sequencing quality check data

| Sample # | Treatment | Conditions | Read 1, before filtering | Read 2, before filtering | Read 1, after filtering | Read 2, after filtering |
|---|---|---|---|---|---|---|
| **1** | 0.1% DMSO | Normoxia | 28059518 | 28059518 | 27705327 | 27705327 |
| **2** | 0.1% DMSO | Normoxia | 51509255 | 51509255 | 50808911 | 50808911 |
| **3** | 0.1% DMSO | Normoxia | 27554576 | 27554576 | 27171360 | 27171360 |
| **4** | 0.1% DMSO | Hypoxia | 28191506 | 28191506 | 27844057 | 27844057 |
| **5** | 0.1% DMSO | Hypoxia | 57091149 | 57091149 | 56317106 | 56317106 |
| **6** | 0.1% DMSO | Hypoxia | 58159615 | 58159615 | 57409120 | 57409120 |
| **7** | 1 μM onametostat | Normoxia | 23317389 | 23317389 | 23027750 | 23027750 |
| **8** | 1 μM onametostat | Normoxia | 53908069 | 53908069 | 53276725 | 53276725 |
| **9** | 1 μM onametostat | Normoxia | 26996981 | 26996981 | 26667743 | 26667743 |
| **10** | 1 μM onametostat | Hypoxia | 26077826 | 26077826 | 25750826 | 25750826 |
| **11** | 1 μM onametostat | Hypoxia | 16846143 | 16846143 | 16759155 | 16759155 |
| **12** | 1 μM onametostat | Hypoxia | 18902202 | 18902202 | 18709772 | 18709772 |

Figure S1. Dynamic range of the resazurin-based viability assay in U-87 MG cells following incubation in normoxia or hypoxia

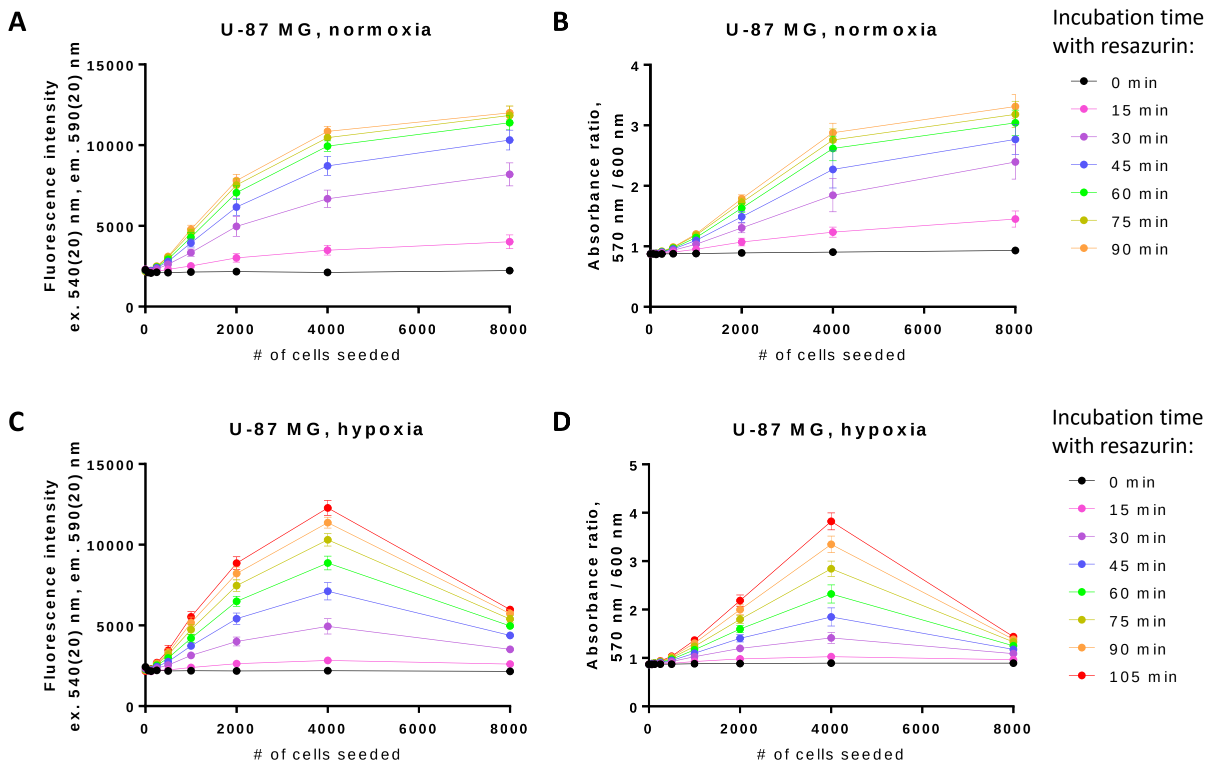


The number of seeded cells is shown on the x-axis. The cells were left to attach for 24 h in normoxia and then were incubated for further 48 h in normoxia (A, B) or in hypoxia (C, D). Graphs A and C feature fluorometric assay format and graphs B and D colorimetric assay format. Data from a single representative experiment is shown; the error bars correspond to the standard deviation of sextuplicates.

Figure S2. Additional radar plots enabling comparison of viability $pIC_{50}$ values obtained in normoxia vs hypoxia

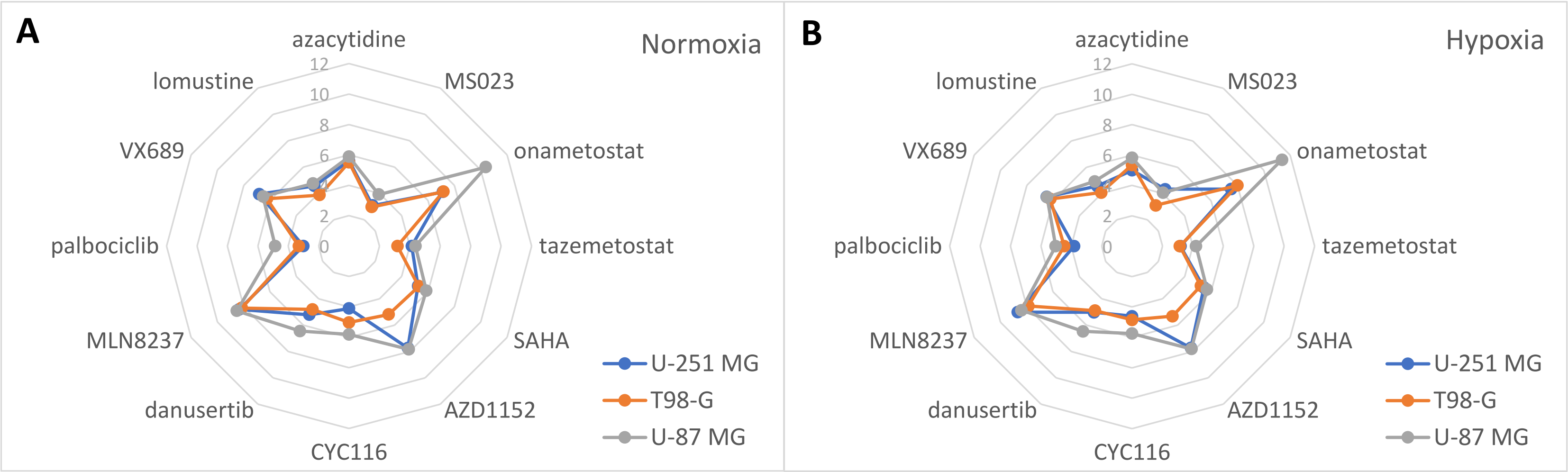


Panel A shows viability profile of different cell lines (legend shown on the right) in normoxia and panel B in hypoxia. The names of compounds are listed along the radar perimeter. The data was obtained by pooling all independent experiments (N ≥ 3). For clarity, no error bars are depicted and only one $pIC_{50}$ value is shown per compound (in case of compounds featuring the biphasic dose-response fit, only the largest $pIC_{50}$ value was chosen). The numbering of y-axis is shown in light grey.

Figure S3. Dose-response curves measured for the compounds of interest in viability assay with T98-G

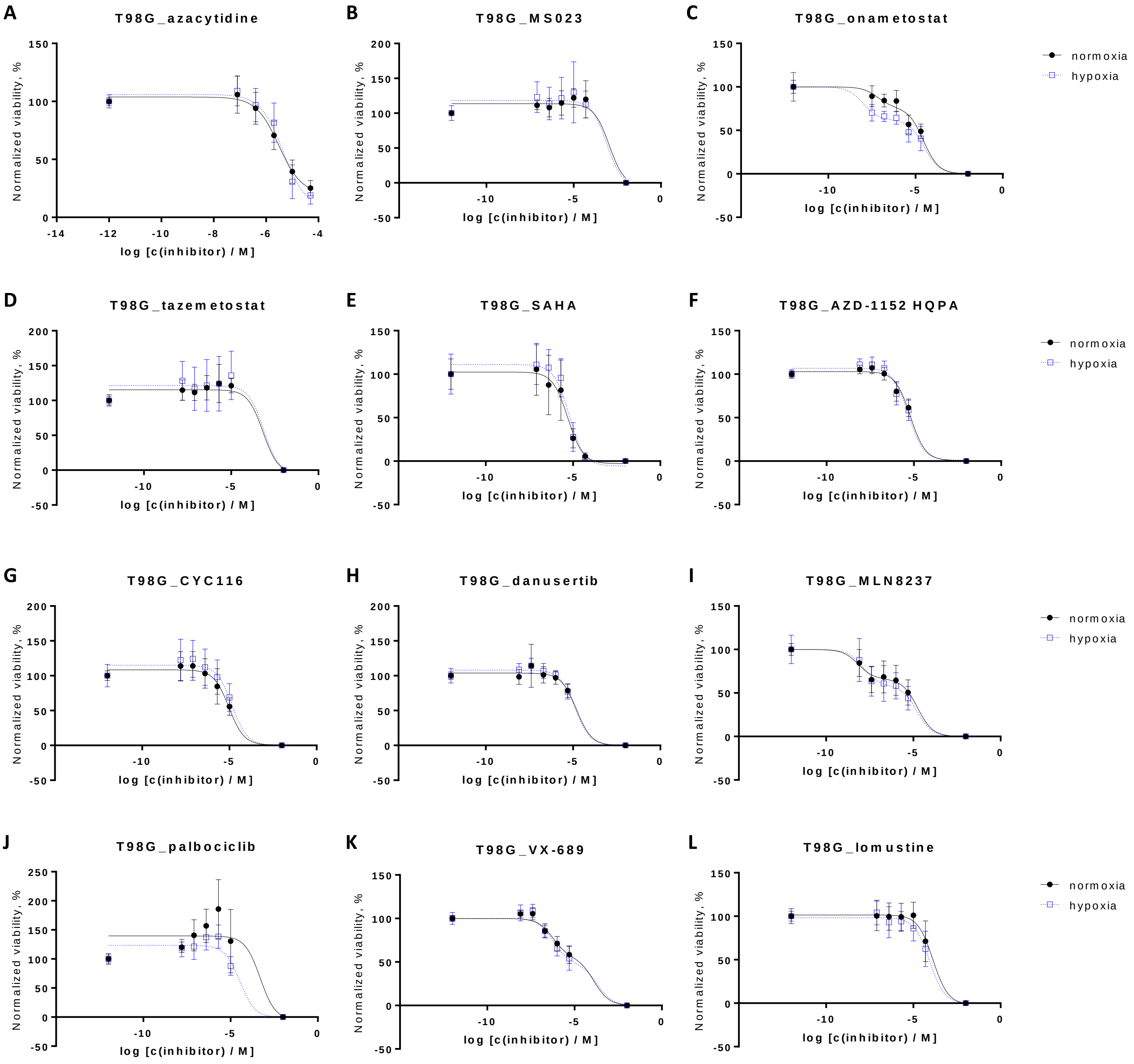


Different panels feature different compounds; the colour code corresponding to the treatment conditions (normoxia or hypoxia) is shown on the right. The left-most point of each dose-response curve corresponds to the negative control (PBS-only treatment) and the right-most point in all panels except A corresponds to the positive control (a well with resazurin but no seeded cells). Pooled normalized data from several independent experiments is shown (N ≥ 3); the error bars indicate the standard deviation. Data points in panels A, B, D-H, J and L were fitted to the logarithmic dose-response function (three parameters), and data points in panels C, I and K to the biphasic equation with the Hill slope values fixed at -1.

Figure S4. Dose-response curves measured for the compounds of interest in viability assay with U-251 MG

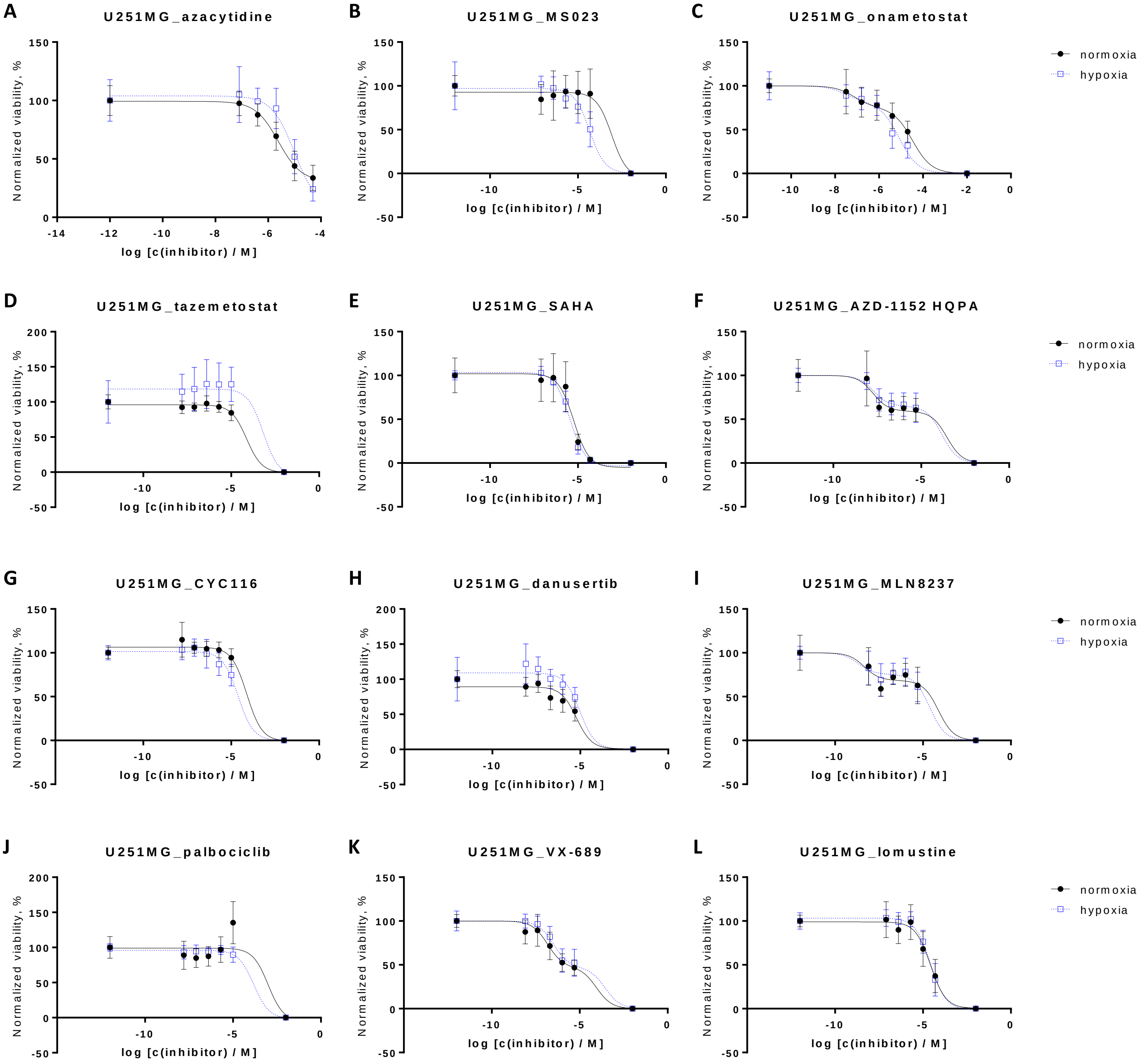


Different panels feature different compounds; the colour code corresponding to the treatment conditions (normoxia or hypoxia) is shown on the right. The left-most point of each dose-response curve corresponds to the negative control (PBS-only treatment) and the right-most point in all panels except A corresponds to the positive control (a well with resazurin but no seeded cells). Pooled normalized data from several independent experiments is shown (N ≥ 3); the error bars indicate the standard deviation. Data points in panels A, B, D, E, G, H, J and L were fitted to the logarithmic dose-response function (three parameters), and data points in panels C, F, I and K to the biphasic equation with the Hill slope values fixed at -1.

Figure S5. Dose-response curves measured for the compounds of interest in viability assay with U-87 MG

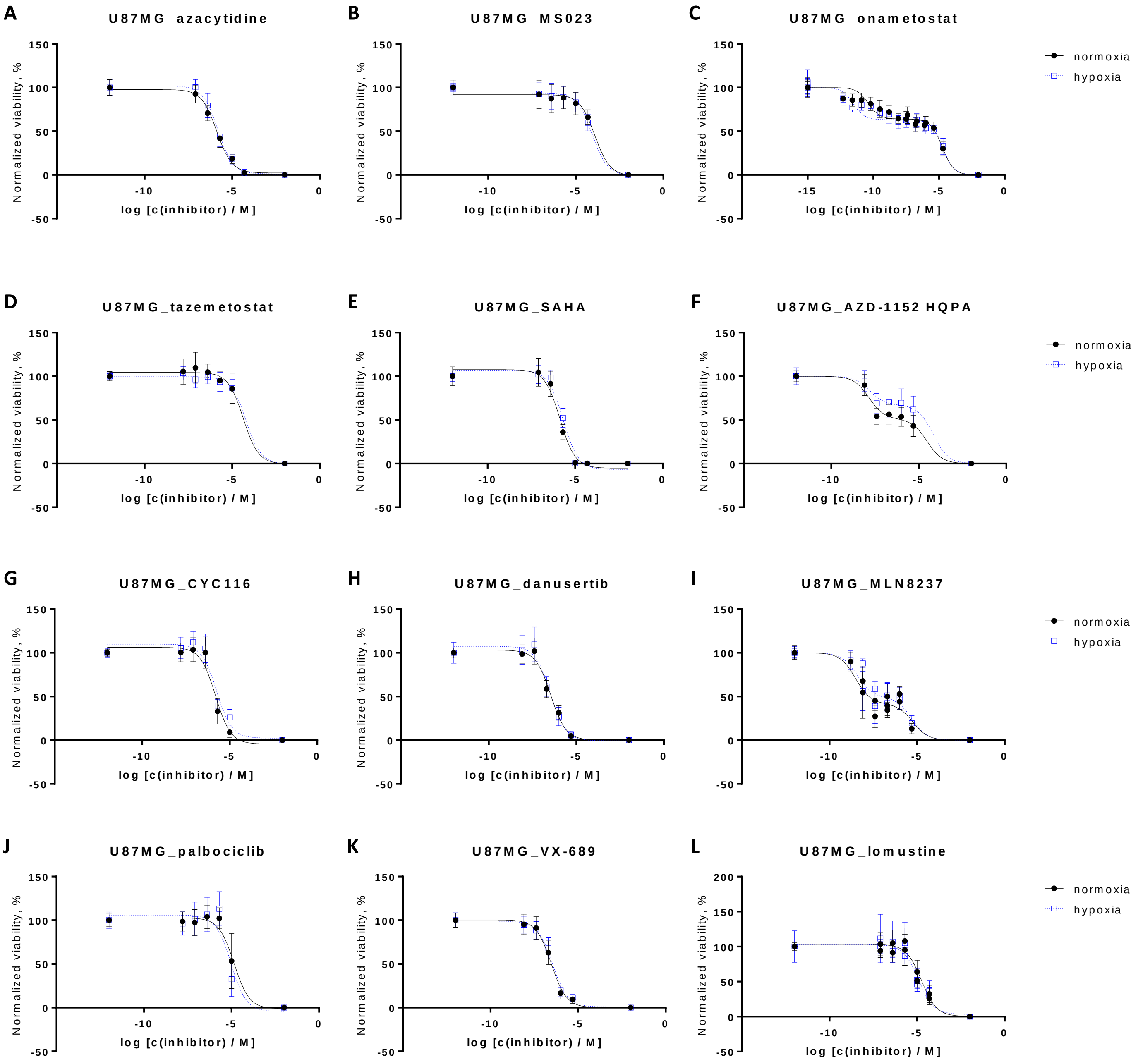


Different panels feature different compounds; the colour code corresponding to the treatment conditions (normoxia or hypoxia) is shown on the right. The left-most point of each dose-response curve corresponds to the negative control (PBS-only treatment) and the right-most point in all panels corresponds to the positive control (a well with resazurin but no seeded cells). Pooled normalized data from several independent experiments is shown (N ≥ 3); the error bars indicate the standard deviation. Data points in panels A, B, D, E, G, H, and J-L were fitted to the logarithmic dose-response function (three parameters), and data points in panels C, F, and I to the biphasic equation with the Hill slope values fixed at -1.

Figure S6. Additional parameters obtained from the dose-response curves in viability assay

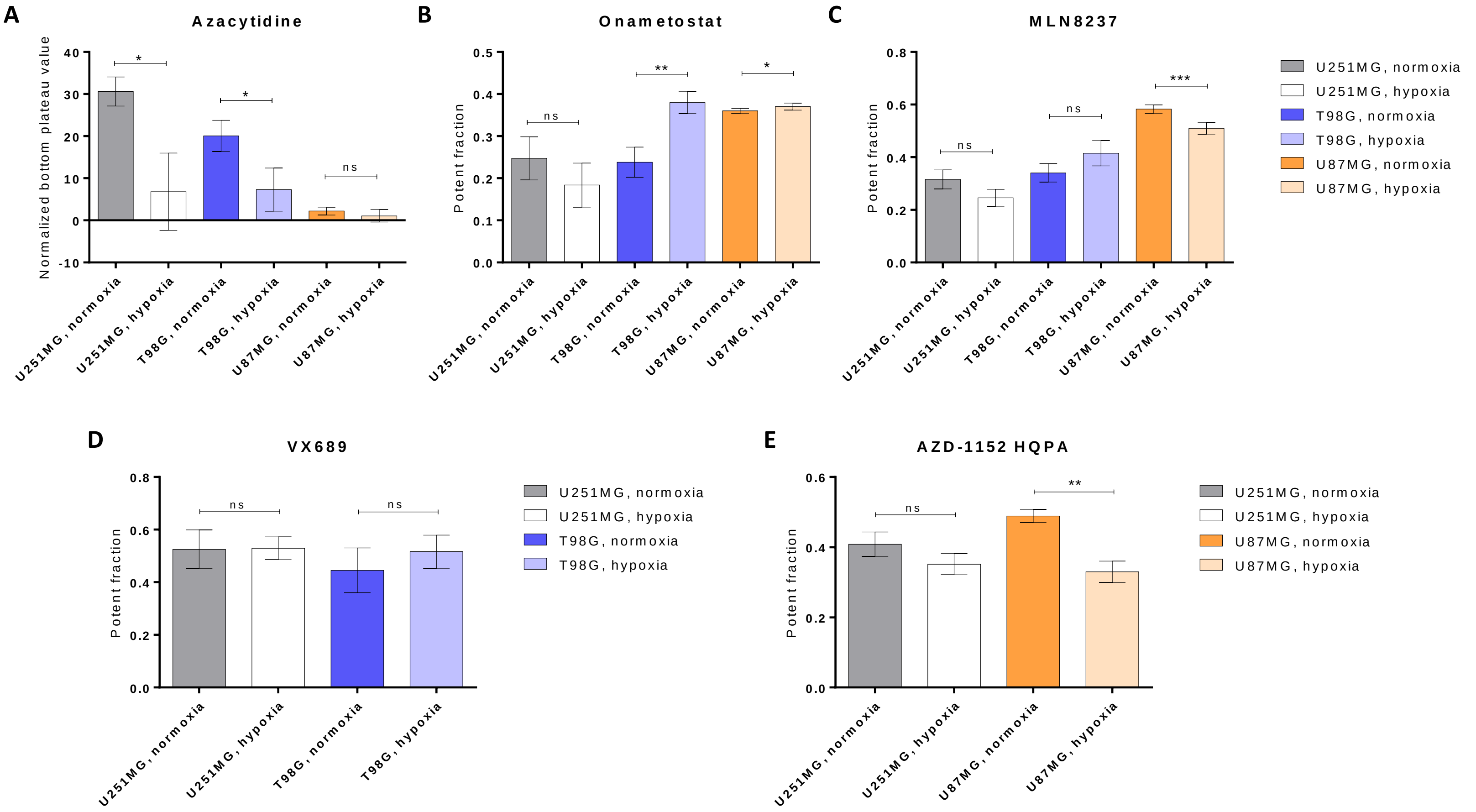


Different panels feature different compounds; the colour code corresponding to the treatment conditions (normoxia or hypoxia) is shown on the right and on the x-axis. Panel A summarizes the bottom plateau values obtained for azacytidine dose-response curves; panels B-E summarize the data on the values of the low-dose fractions in case of compounds featuring biphasic shape of the dose-response curves. Mean values and standard deviations are depicted ($N \geq 3$). The pairwise comparisons show statistical significance of differences for the parameters measured following incubation of cells in normoxia vs hypoxia (unpaired two-tailed t-test with Welch's correction): *** indicates $P \leq 0.001$, ** indicates $P \leq 0.01$, * indicates $P \leq 0.05$, ns indicates not significant.

Figure S7. Propidium iodide staining in glioblastoma cell line spheroids formed in the presence of onametostat or control compounds

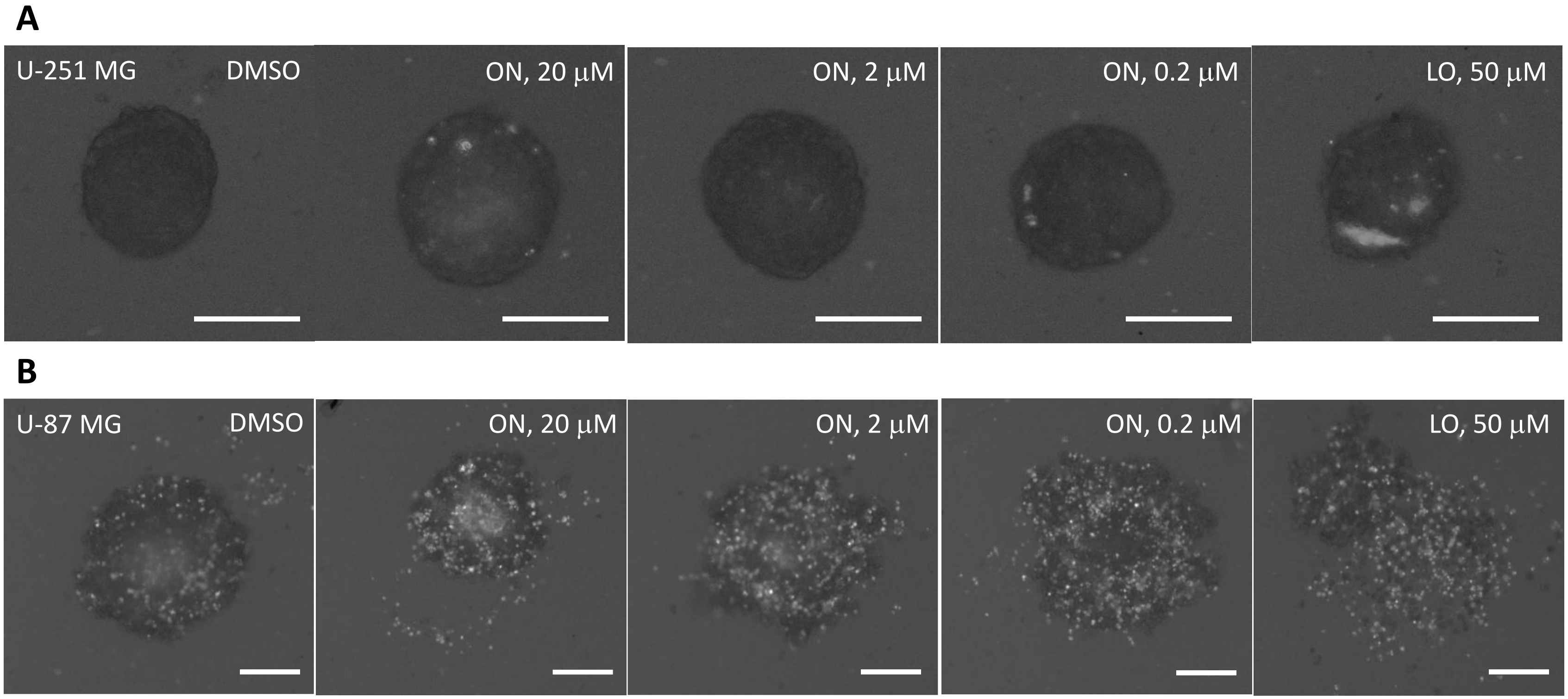


Effect of onametostat (ON) or lomustine (LO) on the spheroid formation in U-251 MG (A) or U-87 MG (B) cells. Panels A and B show examples of spheroid morphology following the 96-h treatment with indicated compounds in a single representative experiment; scale bar: 200 µm. For better visualization, the brightness of all microscopy images was enhanced by 40% and the contrast was reduced by 20%; the quantification of PI signal shown in the main text was done using unmodified images.

Figure S8. Comparison of onametostat effect in normoxic vs hypoxic conditions

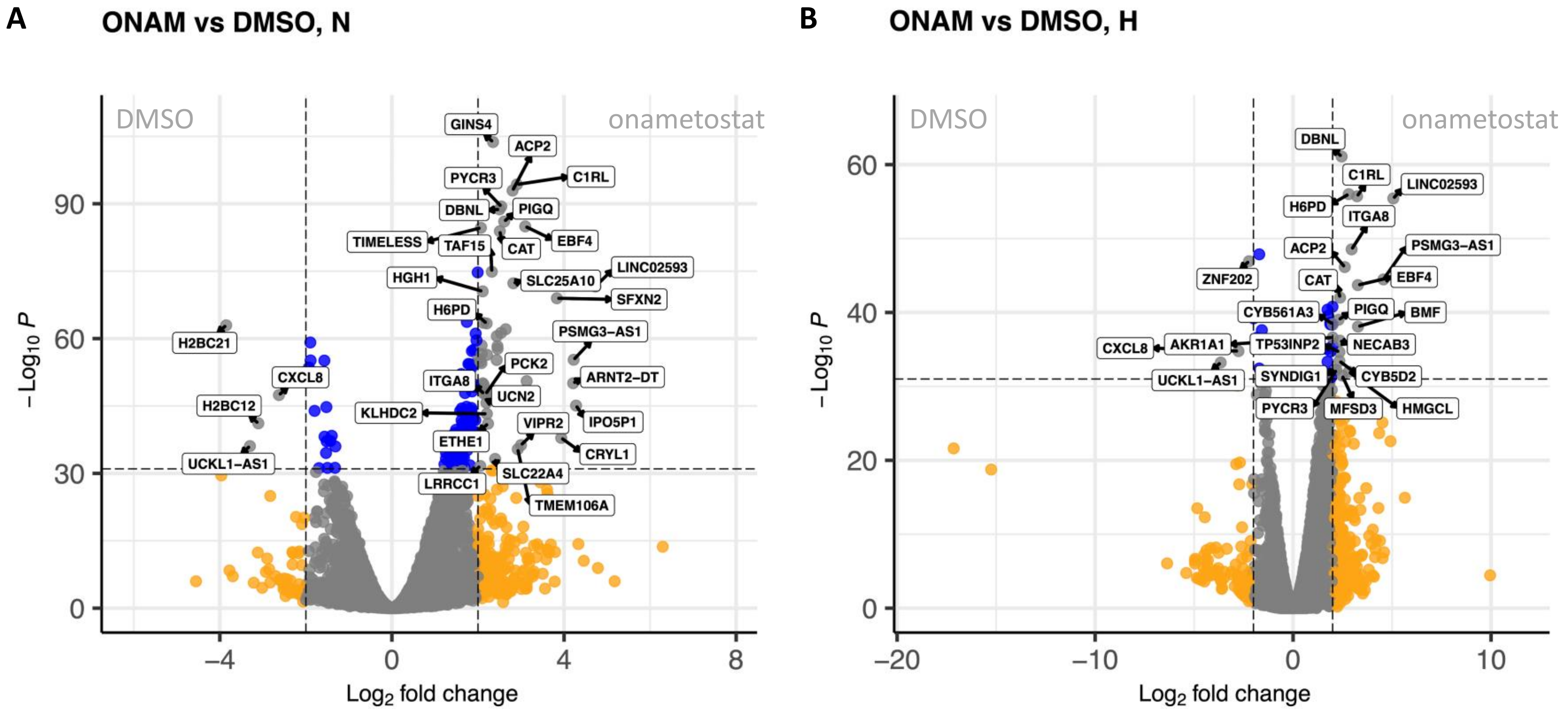


Volcano plots showing DEGs in treatment comparisons. (A) Onametostat- *vs* DMSO-treated T98-G cells following incubation in normoxia. (B) Onametostat- *vs* DMSO-treated T98-G cells following incubation in hypoxia (the panel is identical to the Figure 6B in the main text and is only provided to simplify the comparison). Top hits are marked with the name labels; DEGs coloured in orange possessed binary logarithm of fold change values of below -2 or over 2, while DEGs coloured in blue featured negative logarithm of P-value cut-off of 32. Abbreviations: H, hypoxia; N, normoxia; ONAM, onametostat.
